# A Sensory Processing Hierarchy for Thermal Touch: Thermal Adaptation Occurs Prior to Thermal-Tactile Integration

**DOI:** 10.1101/374447

**Authors:** Hsin-Ni Ho, Hiu Mei Chow, Sayaka Tsunokake, Warrick Roseboom

## Abstract

The brain consistently faces a challenge of whether and how to combine the available information sources to estimate the properties of an object explored by hand. Thermal referral (TR) is a phenomenon that demonstrates how thermal and tactile modalities coordinate to resolve inconsistencies in spatial and thermal information. When the middle three fingers of one hand are thermally stimulated, but only the outer two fingers are heated (or cooled), thermal uniformity is perceived across three fingers. This illusory experience of thermal uniformity in TR compensates for the discontinuity in the thermal sensation across the sites in contact. The neural loci of TR is unclear. While TR reflects the diffuse nature of the thermoceptive system, its similarities to perceptual filling-in and its facilitative role in object perception also suggest that TR might involve inference processes associated with object perception. To clarify the positioning of this thermo-tactile interaction in the sensory processing hierarchy, we used perceptual adaptation and Bayesian decision modelling techniques. Our results indicate that TR adaptation takes place at a peripheral stage where information about temperature inputs are still preserved for each finger, and that the thermal-tactile interaction occurs after this stage. We also show that the temperature integration across three fingers in TR is consistent with precision weighted averaging effect - Bayesian cue combination. Altogether, our findings suggest that for the sensory processing hierarchy of thermal touch, thermal adaptation occurs prior to thermo-tactile integration, which combines thermal and tactile information to give a unified percept to facilitate object recognition.

**Significance Statement:** Thermal touch refers to the perception of temperature of objects in contact with the skin and is key to object recognition based on thermal cues. While object perception is an inference process involving multisensory inputs, thermal referral (TR) is an illusion demonstrating how the brain’s interpretation of object temperature can deviate from physical reality. Here we used TR to explore the processing hierarchy of thermal touch. We show that adaptation of thermal perception occurs prior to integration of thermal information across tactile locations. Further, we show that TR results from simple averaging of thermal sensation across locations. Our results illuminate the flexibility of the processing that underlies thermal-tactile interactions and facilitates object exploration and identification in our complicated natural environment.

## Introduction

Direct manual exploration is regarded as an important and reliable way to obtain information about object properties such as weight and temperature (Lederman and Klatzky, 1997). However, studies have shown that our tactual object perception is not always veridical (Lederman and Jones, 2011). Thermal Referral (TR) is a phenomenon that reflects thermo-tactile interactions in tactual-temperature perception. This illusion was first demonstrated in an experiment wherein three thermal stimulators were touched with the middle three fingers of one hand (from D2 to D4). When the outer two stimulators were warm (cold) and the center stimulator was thermally neutral, warmth (cold) was felt at all three fingers. Notably, this referral of thermal sensation disappeared when the middle finger was withdrawn from the central (neutral) stimulator, indicating that congruent tactile stimulation is essential for TR to occur (Green, 1977).

TR is an important phenomenon that reflects how thermal and tactile modalities coordinate to resolve incoherent spatial information in thermal touch. While it reflects the diffuse nature of the thermoceptive system (Mountcastle, 1961; Han et al., 1998; Bowsher, 2005), its similarities to perceptual filling-in in terms of perceptual continuity and feature averaging (Hsieh and Tse, 2009) and its facilitative role in object perception point to the possibility that TR might involve inference processes related to object perception. Indeed, object inference is important for the everyday activities of recognition, planning, and motor action. It enables us to perceive the surface and material properties of objects quickly and reliably despite the complexity and objective ambiguities in the environments (Kersten et al., 2004). Previous work has shown that inferences regarding visual properties change haptic estimates, such as temperature, weight, surface texture and size (Lederman et al., 1986; Ernst and Banks, 2002; Brayanov and Smith, 2010; Ho et al., 2014). Thus, it is possible that TR involves a cognitive mechanism that assumes homogenous object properties at different points of contact, compensating for the discontinuities in the thermal perception to create a coherent perceptual experience across thermal and tactile modalities.

In this study, we used “perceptual adaptation” as a method to probe the positioning of TR in the sensory processing hierarchy (Heron et al., 2013). Adaptation has been referred to as the psychophysicists’ microelectrode because there is often a strict contingency between adaptation and changes in perception (Frisby, 1979; Thompson and Burr, 2009; Webster, 2011). The perceptual changes that result from adaptation, i.e. the aftereffect, can be used to indicate the sensory coding structure for that dimension. Accordingly, in this study we examined the thermal referral *aftereffect* in order to elucidate the perceptual coding structure that underlies TR in particular, but thermal-tactile interactions more generally. We compared the perceptual aftereffects when the participants were adapted to the classical 3-finger configuration of TR, or a perceptually equivalent uniform stimulation. We found that when asked to report their thermal perception across all three fingers together, the induced perceptual aftereffect was equivalent to that following adaptation to a truly uniform thermal display. However, when asked to report about their thermal perception for only a single finger at a time, participants reports were instead consistent with having adapted to the physically presented stimulation. To resolve these apparently inconsistent results, in a second experiment we used a precision-weighted Bayesian cue combination model to investigate how thermal information from each finger contributed to global perception across the hand under TR. Our results indicated that TR is consistent with Bayesian combination of the thermal inputs from each finger in the hand and that the results from the first experiment reflect a process that involves only peripheral adaptation. Overall, our results demonstrate that thermal adaptation occurs prior to thermo-tactile integration and that the human brain uses strategies common to other sensory dimensions to infer the combined thermal properties across the hand. These findings support the idea that the combination of different features in thermal-tactile perception follows inference processes for the purpose of coherent object perception.

## Materials and Methods

### Participants

Eleven naıve paid volunteers (three males and eight females) and three authors HH (female), DC (female) and WR (male) participated in Experiment 1. Eleven naïve paid volunteers (two males and nine females) and two authors HH (female) and ST (female) participated in Experiment 2. Five participants, including the author HH, were common to both experiments. All the participants were right-handed, except ST in Experiment 2. The participants aged between 22 and 45 years, and had no known abnormalities of their tactile and thermal sensory systems. Recruitment of participants and experimental procedures were conducted in accordance with the *Declaration of Helsinki*.

### Apparatus

To adapt both participants’ hands to a preset temperature, two thermal displays – hereafter, termed adapting thermal displays - were used. A custom made hot plate (180 × 180mm) consisting of heating wire (Yagami Inc, Nagoya, Japan) and copper plate was used for reference (dominant) hand adaptation, and a thermal display made of three copper bars, two of which controlled by a water-heating/cooling system (Eyela NCB-1200, Tokyo Rikakikai Co. Ltd, Tokyo, Japan) and one of which by electric heater (Takagi Mfg. Co., Ltd., Tsukuba, Japan), was used for test (non-dominant) hand adaptation. The two adapting thermal displays were put near each hand respectively (Figure 1). In between the adapting thermal displays, another two thermal displays –hereafter, termed testing thermal displays - were used to present test thermal stimulation to the middle three fingers of each hand (Figure 1). Each thermal display consisted of three Peltier devices with a surface area of 20 × 20 mm (FPH1-7106M, Fujitaka Co., Kyoto, Japan). The Peltier devices were housed in plastic holders, which expose a constant surface area of 300 mm^2^ of the Peltier devices to the participant’s fingerpad. Two digital–analog converters (ADI16-16 and DA16-16, Contect Co., Osaka, Japan) and a PI control loop programmed in Matlab (Mathworks, Inc., MA, USA) were employed to control the surface temperatures of the Peltier devices. The temperature feedback was provided by thermistors (457 *μ*m in diameter and 3.18 mm in length; 56A1002-C8, Alpha Technics, CA) sandwiched between the Peltier devices and plastic holders. The maximum rate of temperature change was 10°C/sec for cooling and 18°C/sec for heating. Achieving a steady state took about 1 sec. After a steady state had been reached, the temperature of each Peltier device could be maintained within 0.5°C of the desired temperature. To facilitate heat dissipation, the testing thermal displays were placed on top of copper heat sinks (P-200S, Takagi Mfg. Co., Tsukuba, Japan) connected to a water-cooling system (Eyela NCB-1200, Tokyo Rikakikai Co. Ltd, Tokyo, Japan).

**Figure 1.**
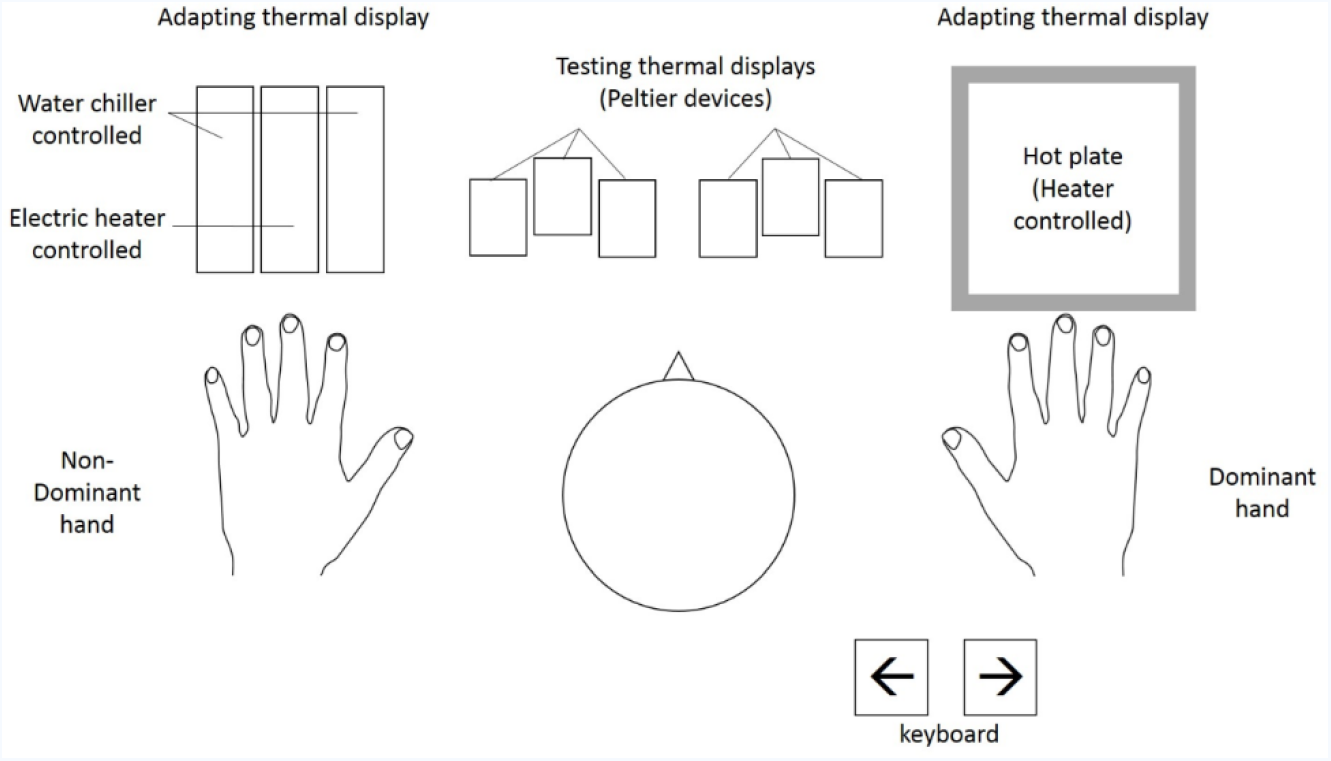
Experimental thermal apparatus set up for this study. In the experiments we used a two-hand configuration. The dominant hand touched the reference stimulation and the non-dominant hand touched the test stimulation. The adapting thermal displays at two sides were used to adapt both participants’ hands to a preset temperature. After adaptation, the participants moved their hands to the testing thermal displays to feel the test thermal stimulation applied to the middle three fingers of each hand. The participants reported which hand (left or right) felt warmer by pressing left or right arrow keys of the keyboard.

### Experiment 1: Adaptation to illusory thermal uniformity

As there are individual differences in experiencing the thermal referral illusion (Ho et al., 2011), before the main experiment, each participant completed a matching experiment. This experiment aimed to find the temperature D2 and D4 should touch (*T*°C) such that, when presented in combination with the middle finger touching neutral temperature at 33°C, a thermal referral percept that is perceptually indistinguishable from a uniform presentation of 35°C across all fingers was generated. Note that 35 °C was chosen because (1) it is warm given a baseline temperature of 33 °C; (2) it can be fully adapted within the initial adaptation duration of 10 minutes (Kenshalo and Scott, 1966); and (3) the outer finger temperature (*T*°C) that needed to induce an illusory thermal uniformity of 35 °C is reported to be about 38-39 °C (Ho et al., 2011), which is below pain threshold (Greenspan and Kenshalo, 1985). At the beginning of the experiment, participants placed both of their hands on the adapting thermal displays which were set at the neutral temperature of 33°C. Upon hearing a sound cue, participants moved their hands to the test thermal displays. Their reference (dominant) hand touched the uniform temperature at 35°C, and test (non-dominant) hand touched the three-stimulator thermal display, in which the central stimulator was set at 33°C and the temperature of the outer stimulator varied between 35, 37, 39, 41, 43°C between trials. Another sound cue was presented after 5s to cue participants to lift both hands off the thermal display and report which hand (left or right) felt warmer by pressing left and right arrow keys of the keyboard. Each stimulus temperature was repeated 12 times, giving a block of 60 trials that were presented in randomized order. There were three blocks in this matching experiment, which gave a total of 180 trials. Each block of trials lasted for about 20 min, and there was at least a 20-min break between the blocks. Each participant’s data were fitted with a cumulative Gaussian function, to estimate the Point of Subjective Equality (PSE) and the standard deviation of the participants’ responses (for details of the fitting, see Experimental Design and Statistical Analysis section). The PSE told us about the temperature combination of thermal referral stimulation that would be equivalent for a physical uniform temperature 35°C for each participant. The outer finger temperature (*T*°C) was found to be in the range of 35.8 −38.7 °C, with a mean of 37.1 °C. These values were used in subsequent experiments.

In the main adaptation experiment, the aftereffect that we were looking at was the “new physiological zero” created after adaptation. Physiological zero refers to the temperature that feels thermally neutral. As there is no fixed reference point for temperature perception, one’s “physiological zero” can be manipulated through adaptation (Stevens, 1991). The subsequent thermal perception (aftereffect) is referenced to this new physiological zero and a temperature above (below) which would be felt as warm (cold). A famous example is the “three-bowl illusion”, wherein 27°C water can feel either warm or cold depending on to which temperature the hand previously adapted (Tritsch, 1988). Accordingly, in this experiment, we have participants adapted to different patterns of temperature stimulation, including the classical 3-finger configuration of TR stimulation. After adaptation, participants touched a test stimulus and reported the thermal sensation elicited, i.e. the aftereffect. From these data, we were able to estimate the “new physiological zero” for different adaptation conditions.

To look at the effect as a whole and in relation to each individual finger, we manipulated the finger configuration in the adaptation and test phases. In the whole hand (3-finger) condition (Figure 2A), the test hand (non-dominant hand) of participants was adapted to physically uniform stimulation and TR stimulation in different sessions. When adapting to physically uniform stimulation, participants touched the adapting thermal display which were all set to 35°C. When adapting to TR stimulation, participants touched the adapting thermal display in which the temperature of the center stimulator was set to 33°C, and the two outer stimulators were set at *T*°C (which was the temperature matched for each participant before the adaptation experiments). At the same time, the reference hand (dominant hand) of participants touched the hot plate set at 35°C. The initial adaptation duration at the beginning of each session was 10 minutes to ensure adequate exposure to the adapting stimuli. Upon hearing a sound cue, participants moved all three fingers of both hands and touched the testing thermal displays. In this test phase, the test hand touched test stimuli varied between 31, 33, 35, 37, 39, 41°C (same across three channels), and for the reference hand always 35°C to create neutral thermal sensation. Another beep sound (after 5s) was presented when participants lifted their hands off the testing thermal displays and used the arrow keys on the keyboard to indicate which hand (left or right) feels warmer. Each stimulus temperature was repeated 12 times, giving a block of 60 trials that were presented in randomized order. There were three blocks in each experimental session, giving a total of 180 trials in each session. Each block of trials lasted for about 30 min, and there was at least a 30-min break between the blocks. Each participant’s data were fitted with a cumulative Gaussian function, to estimate a Point of Subjective Equality (PSE) and the standard deviation of the participants’. Given that the reference hand always felt thermally neutral, the PSE gives the temperature that felt thermally neutral to the test hand, which in turn corresponded to the new physiological zero after adaptation.

**Figure 2.**
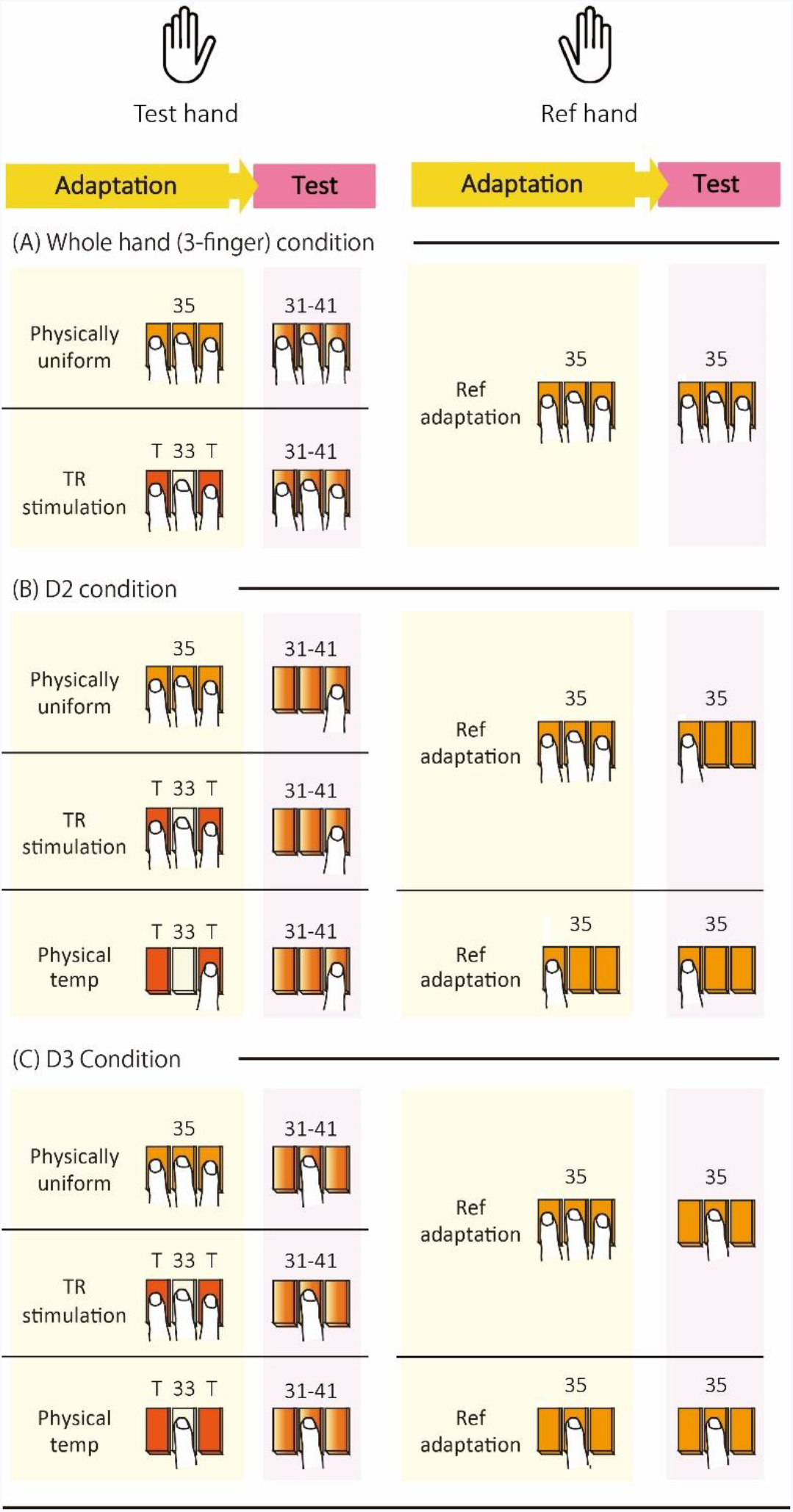
Experimental conditions in Experiment 1. In this experiment, each participant’s test hand was adapted to different patterns of temperature stimulation, including the classical 3-finger configuration of TR stimulation (TR stimulation), matched physically uniform stimulation (physically uniform) and physical temperatures presented in TR stimulation (physical temp). At the same time, the participant’s reference hand was adapted to the reference temperature of 35°C. After adaptation, the participants touched a test stimulus varied between 31-41°C with their test hand and a test stimulus set at the reference temperature of 35°C with their reference hand. The participant’s task was to report which hand felt warmer. From these data, we were able to estimate the “new physiological zero” for different adaptation conditions. To look at the effect as a whole and in relation to each individual finger, we manipulated the finger configuration in the adaptation and test phases. (A) The whole hand (3-finger) condition, wherein the middle three fingers of one hand were used in both adaptation and test phases. (B) D2 condition, wherein only D2 was used in the test phase. In the adaptation phase, the middle three fingers of one hand were used for physically uniform stimulation and TR stimulation and only D2 was used in physical temperature stimulation. (C) D3 condition, wherein only D3 was used in the test phase. In the adaptation phase, the middle three fingers of one hand were used for physically uniform stimulation and TR stimulation and only D2 was used in physical temperature stimulation.

To look at the effect for each individual finger, we also tested D2 and D3 separately after adapting to three different temperature patterns – Physically uniform stimulation, TR stimulation and physical temperature (Figure 2B & 2C). The procedure of adapting to physically uniform stimulation and TR stimulation were identical to that of the three-finger condition, except that at the test phase, only one of the fingers (D2 or D3) of both hands touched the test thermal display (Figure 2B & C). In the physical temperature adaptation, D2 and D3 adapted to their corresponding physical temperature separately (D2 to *T* °C and D3 to 33 °C). The procedure was similar to that of the other two conditions, except that at both adaptation and test phase, only one of the fingers (D2 or D3) of both hands touched the thermal displays. The thermal perception of D2 or D3 was tested individually in separate sessions and participants were told which finger was the test finger at the beginning of each session. Note that among the three adaptation patterns used, physically uniform stimulation and the physical temperature represent two bounds of possible outcomes of TR stimulation adaptation, with the physically uniform stimulation giving the aftereffect more like having adapted to the perceptual value of TR (i.e. illusion) and the physical temperature giving the aftereffect more like having adapted to the physical values of TR.

### Experiment 2: Combination of thermal information across fingers

The aim of this experiment was to understand how the thermal inputs from each finger was combined to reach a final percept global percept of uniformity seen in thermal referral. Precision-weighted (Bayesian) averaging, in which the independent inputs are combined based on a weighting proportional to the inverse of their variability, is a common approach used to describe how different sources of perceptual information are combined (Ernst and Banks, 2002; Alais and Burr, 2004; Körding et al., 2007; Shams and Beierholm, 2010). To conduct this analysis, we measured the thermal perception, and variability in thermal perception, for each finger individually and altogether under TR stimulation. We then compared the obtained results for the TR condition with what we would predict under a precision-weighted model based on the thermal perception for each finger alone.

In this experiment, the TR stimulation was set at [37, 33, 37]°C for D2, D3 and D4, respectively. 37°C was used for the outer two fingers because it was the average value found in the matching experiment. The sensory variability of D2, D3 and D4 and three fingers all together were measured in separate experimental sessions, conducted in a pseudo-randomized order. At the beginning of each experimental session, both hands were initially adapted to neutral temperature of 33°C. Upon hearing a sound cue, participants moved their hands to the test thermal displays. When testing individual fingers (D2, D3 or D4), the finger of the reference (dominant) hand touched the corresponding temperature stimulation (D2 and D4 to 37°C, D3 to 33°C), and the finger of the test (non-dominant) hand touched the three-stimulator thermal display, in which the temperature of the corresponding stimulator varied between 33-41°C for D2 and D4 and 29-37°C for D3 between trials (Figure 3). Another sound cue was presented after 5s to cue participants to lift both hands off the thermal display and report which hand (left or right) feels warmer by pressing left and right arrow keys of the keyboard. When testing three fingers all together, the reference (dominant) hand touched the TR stimulation, and the test (non-dominant) hand touched the three-stimulator thermal display, in which the temperatures varied between 31-39°C (same across 3 channels) between trials (Figure 3A). For all four conditions, each stimulus temperature was repeated 12 times, giving blocks of 60 trials that were presented in randomized order. There were three blocks in each experimental session, giving a total of 180 trials for each session. Each block of trials lasted for about 30 min, and there was at least a 30-min break between the blocks. Each participant’s data were fitted with a cumulative Gaussian function, to estimate a Point of Subjective Equality (PSE) and the standard deviation of the participants’ responses. The PSE told us the apparent intensity of TR stimulation and the standard deviation informed us the sensory variability in the estimation (for details of the fitting, see Experimental Design and Statistical Analysis section).

**Figure 3.**
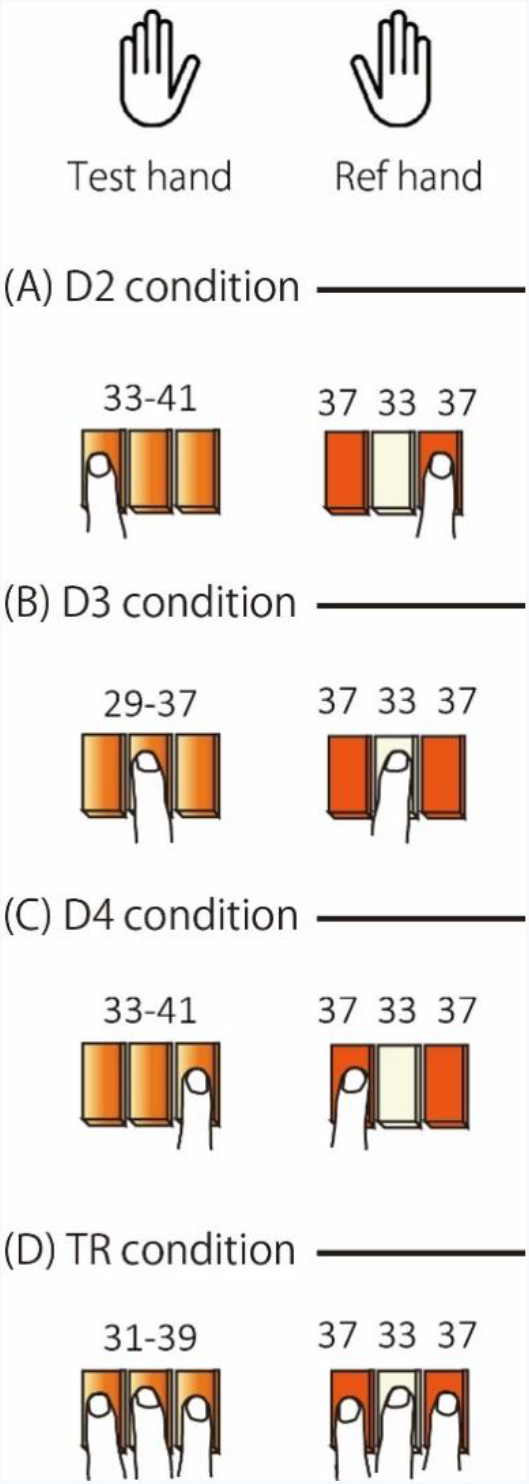
Experimental conditions of Experiment 2. In this experiment, we measured the sensory variability of the fingers individually and altogether under TR stimulation. We used two hand configuration and both hands were initially adapted to the neutral temperature of 33°C. In the test phase, the test hand touched a thermal stimulus that varied in a range between 29-41 °C and the reference hand touched the TR stimulation. We asked participants to compare the thermal sensations between two hands for each finger individually (A-C), or in combination for thermal stimulation that would produce TR illusion (D).

### Experimental Design and Statistical Analysis

In all conditions in Experiments 1 & 2, the test and reference hands were tested simultaneously and a single interval forced choice procedure was adopted, in which the participants were instructed to report which hand (right or left) felt warmer. In each condition, the proportion of ‘warmer’ responses for the five levels of stimulus temperature were fitted with a cumulative Gaussian function. The two parameters, the mean and the standard deviation, were obtained by a maximum likelihood procedure. The mean of the fitted function was used as an estimate for the Point of Subjective Equality (PSE) between two hands while the standard deviation indicates the precision with which the participant made the categorization of relative warmness (coolness).

The standard deviation for the fitted cumulative Gaussian was used as a criterion to exclude participants from further analysis in the cases where the participant performance on the task was too poor for the results to be of any meaning. Participants who had a standard deviation more than three times that of the group mean were excluded from further analysis because this high variability in the responses indicated that they could not easily discriminate which hand was warmer for the range of thermal differences presented. Following this exclusion criteria, 10 participants in Experiment 1 and 11 participants in Experiment 2 remained.

Bayesian statistics were used to analyze the data from Experiments 1 & 2 with JASP (JASPTeam, 2018). In Bayesian tests, the Bayes factor, BF_10_, indicated the relative strength of evidence for an alternative hypothesis to the null hypothesis. More specifically, a Bayes factor of *x* indicates that the data are *x* times more likely under the alternative than under the null hypothesis. The value of Bayes factor is continuous and varies between 0 and ∞. A Bayes factor of 1/3 or less is commonly taken as evidence for the null hypothesis and of 3 or more as substantial evidence against the null (Dienes, 2014). Bayes factors are in particular useful when non-significant results are obtained with traditional Frequentist tests because they provide an approach for discriminating whether non-significant results support a null hypothesis over a theory or whether the data are just insensitive.

## RESULTS

### Experiment 1: Adaptation to illusory thermal uniformity

Here we show that when observers reported about all three fingers together, the PSE following TR adaptation (Test 2: mean PSE= 35.0 °C) was nearly identical to that following adaptation to the subjectively matched physical thermal uniformity (Test 1: mean PSE= 35.0 °C), as shown by the Bayesian Paired Samples T-Test (BF_10_ = 0.31 see Figure 4). This result indicates that adapting to TR produces an aftereffect commensurate with the perceived, rather than physical temperatures of the fingers. However, when reporting each finger in isolation, a different pattern of results is seen. Here, the data show strong evidence that the PSE following the TR adaption (Test 4 for D2: PSE= 35.3 °C, or Test 7 for D3: PSE= 33.5 °C) was different from that of the subjectively matched physical thermal uniformity (Test 3 for D2: PSE= 34.6 °C or Test 6 for D3: PSE= 34.6 °C), as shown by Bayesian Paired Samples T-Tests (D2: BF_10_ = 22.01; D3: BF_10_ = 3.89; see Figure 4). Additionally, our data indicate a trend towards the PSE following TR adaptation (Test 4 for D2: PSE= 35.3 °C, and Test 7 for D3: PSE= 33.5 °C) being the same as having adapted to the local finger by itself (Test 5 for D2: PSE= 35.6 °C and test 8 for D3: PSE= 33.4 °C). However, the evidence here is not strong enough to accept or reject the null hypothesis that the PSE of these two conditions are the same as shown by the Bayesian Paired Samples T-Test (D2: BF_10_ = 0.71; D3: BF_10_ = 0.40; see Figure 4).

**Figure 4.**
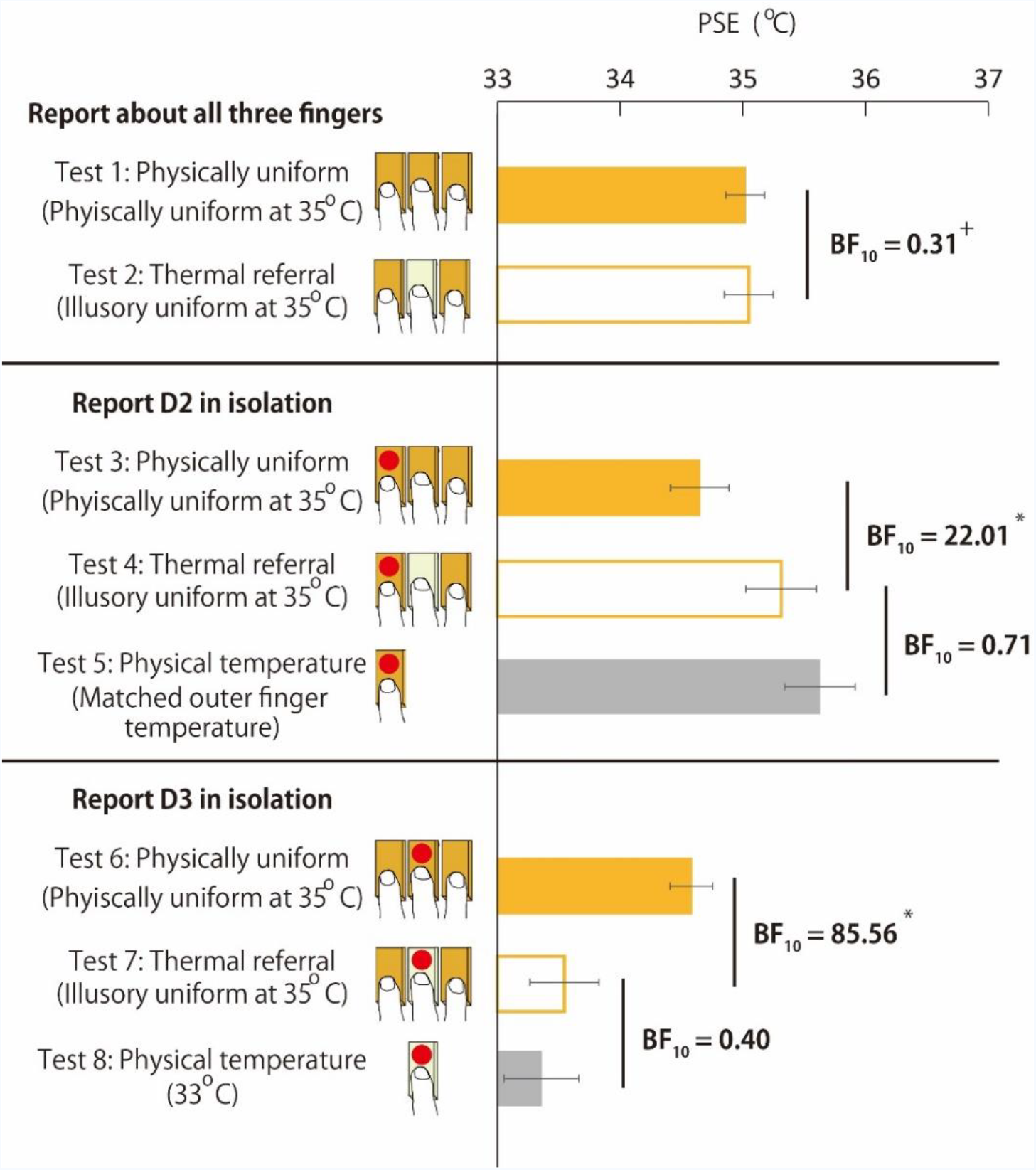
Aftereffects of adapting to different thermal stimulation patterns (TR, perceptually equivalent physically uniform stimulation, or each finger stimulated in isolation) under different configurations of fingers during testing (All fingers, D2, or D3 in isolation). Errorbars indicate standard error of means. The asterisk indicates substantial evidence against the null hypothesis (BF_10_ >3) and the plus sign indicates substantial evidence for the null hypothesis (BF_10_ < 1/3).

Overall, these results indicate that the aftereffect depended on the spatial configuration of the fingers in the test phase. When asking about the thermal perception of the whole hand (3-finger) participants’ responses were like having adapted to the illusory rather than physical stimulation. In contrast, when asking about the individual finger perception, participants’ reports were more like having adapted to the physical values. One possible explanation is that adaptation effect depends on spatial contingency of the fingers. Similar to the McCollough effect, wherein color aftereffects are contingent on the presence of particular grating orientation (McCollough, 1965), only in the presence of the correct spatial configuration of fingers (all three in contact with the thermal surface), we obtain a pattern of results consistent with the illusion adaptation. This explanation in turn implies that the physiological zero for tactual temperature estimation would depend on the consistency of finger configuration to a recent sensory history. This is, however, highly speculative as the resetting of the physiological zero for temperature perception has been shown to be a rather automatic process, relating to change in the temperature threshold for TRPM8 activation and responses in DRG neurons (Fujita et al., 2013). In line with this view, our pattern of results may instead reflect a process that involves only peripheral adaptation, with adaptation to the illusory TR never occurring. Following only peripheral adaptation, such that each finger was adapted to a new physiological zero, presenting the 3-finger configuration in the test phase with a uniformly thermal test display would lead to different apparent temperatures across the 3 fingers. This inhomogeneous temperature distribution could lead to another TR-like temperature redistribution (Green, 1977) – effectively the reverse TR to that presented in the adaptation period, with the thermal sensation for D2 and D4 being that of coolness, and the sensation for D3 being warm. If the sum of these thermal sensations is similar to 35°C, we would find a matched PSE (see Test 2, see Figure 4). This explanation is based on the assumption that the whole hand estimate results from some kind of simple combination of the thermal estimates produced for each finger. To test this assumption, Experiment 2 was designed to investigate how thermal information across fingers are combined to reach final percept.

### Experiment 2: Combination of thermal information across fingers

In this experiment, we measured perception of thermal sensation for D2, D3 and D4 individually and altogether for thermal inputs that would produce TR. The PSEs and SDs are listed in Table 1. The results of PSEs agree with the previous findings that the illusory thermal sensation perceived at D3 is not simply a “copy” of the thermal changes applied to D2 and D4 (Ho et al., 2011; Cataldo et al., 2016). This is because, if it were the case, the PSE of TR would be similar to those of D2 and D4. The results listed in Table 1 show clearly that this was not the case, as the PSEs of TR were *lower* than those of D2 and D4 and *higher* than that of D3 for all participants, indicating an averaging process is involved to combine the thermal changes applied to the three fingers. The combination of different sensory estimates according to a precision-weighted (Bayesian) scheme is a commonly found strategy in human perception (Ernst and Banks, 2002; Alais and Burr, 2004; Shams and Beierholm, 2010). In the precision-weighted model, the independent estimates of D2, D3, and D4, *Ŝ*_D2_, *Ŝ*_*D*3_ and *Ŝ*_*D*4_, i.e. the PSEs of D2, D3, and D4, are combined based on a weighting proportional to the inverse of their variability (Alais and Burr, 2004):

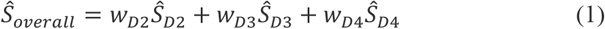

where *w_D2_* is inversely proportional to their variances:

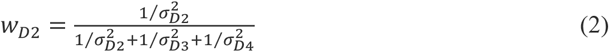

and likewise for *w_D3_* and *w_D4_*. Accordingly, the less variable the estimate, the higher the weighting would be in combination. In other words, when combining multiple estimates of the same property, more trust is given to the more reliable information source.

**Table 1.** PSEs and SDs for temperature estimation at each finger and 3 fingers as a whole under TR stimulation. Shaded participants are those whose SD for measured TR was smaller than the smallest SD of D2, D3 or D4. Units in degree Celsius.

| Participant | SD |  |  |  | PSE |  |  |  |
| --- | --- | --- | --- | --- | --- | --- | --- | --- |
|  | D2 | D3 | D4 | TR | D2<br>(37) | D3<br>(33) | D4<br>(37) | TR |
| 1 | 1.5 | 2.3 | 1 | 1 | 37.9 | 33.5 | 38.1 | 36.3 |
| 2 | 2 | 0.9 | 4.2 | 1.6 | 35.5 | 32 | 37.7 | 34.5 |
| 3 | 1.1 | 0.3 | 0.4 | 0.9 | 36.6 | 32.7 | 37.1 | 36.3 |
| 4 | 3.3 | 1.3 | 0.3 | 1.1 | 35.7 | 30.7 | 35.1 | 34.3 |
| 5 | 3.1 | 2.4 | 2.8 | 1.8 | 39.4 | 32.7 | 38.3 | 37 |
| 6 | 1.5 | 1 | 1.4 | 1.3 | 38.1 | 32.4 | 36.9 | 36 |
| 7 | 1.4 | 1.3 | 2 | 1.1 | 36.3 | 33.7 | 36.9 | 34.9 |
| 8 | 0.2 | 1 | 1.7 | 0.4 | 37.1 | 32.8 | 39.7 | 35.1 |
| 9 | 0.9 | 1 | 1 | 0.2 | 35.9 | 31.5 | 37.2 | 35.9 |
| 10 | 1.7 | 1.7 | 3.3 | 0.7 | 36.8 | 32.6 | 38.8 | 35.6 |
| 11 | 1.2 | 1.1 | 2 | 1.1 | 37.1 | 34 | 38.3 | 35.8 |
| mean±SE | 1.6±0.3 | 1.3±0.2 | 1.8±0.4 | 1.0±0.1 | 36.9±0.4 | 32.6±0.3 | 37.6±0.4 | 35.6±0.2 |

Looking at the performance of the precision weighted model estimates, based on the single finger participant estimates, we see that the Bayesian approach does a good job of describing the participants’ reports under the three-finger TR condition (see Figure 5, Bayesian Paired Samples T-Test: BF_10_ = 0.37). While the Bayesian model estimates are broadly consistent with the obtained TR results, we also wanted to check its performance against the simplest possible strategy available in this case – a simple linear average. Again, as seen in Figure 5, the linear averaging also produced estimates largely consistent with participants’ reports under TR (Bayesian Paired Samples T-Test: BF_10_ = 0.34). These comparisons between the measured TR to the estimates produced by precision weighted model and linear averaging model did not provide strong support for either precision weighted or linear estimates over each other. That our data do not strongly support one combination rule or the other is probably due to the fact that the precision of temperature estimation was similar among three fingers – conditions that approximate linear averaging under the precision weighted scheme. A strong prediction that might differentiate the Bayesian from linear averaging approaches is that the precision-weighted model also specifies that the overall precision of the combined estimate is better than the best precision of any single contributing estimate (i.e. the variability of the estimate in the TR condition should be lower than that obtained in any of the single finger conditions). Examining participants’ reports, we find evidence that this is true when looking at the group mean SD (TR: Mean SD = 1.0; D2: mean SD = 1.6; D3 = mean SD= 1.3; D4: mean SD = 1.8; see Table 1); however, when looking at the individual data, only 4/11 of the participants fulfilled this requirement (shaded participants in Table 1). Overall, these results indicate that a simple averaging scheme (either Bayesian or linear) is used for the integration of thermal information across three fingers to produce the TR illusion.

**Figure 5.**
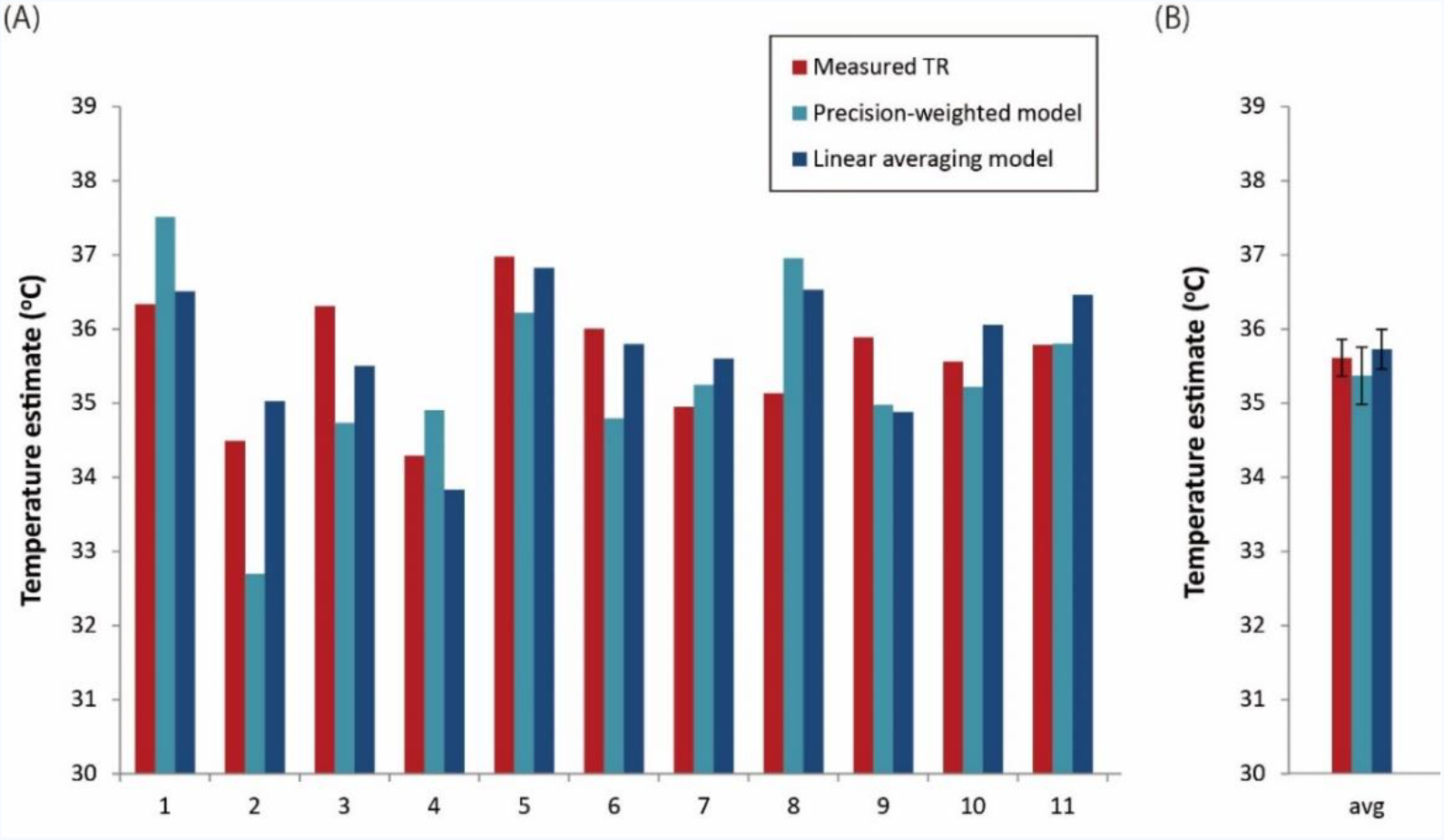
Comparison of actual and predicted apparent temperatures for TR stimulation. (A) Measured TR (red bars), the precision weighted model estimates (light blue bars) and the linear averaging model estimates (dark blue bars) for 11 participants. The measured TR is the Point of Subjective Equality (PSE) obtained in TR condition (see Figure 3D). The precision weighted model estimates were calculated according to Equations 1 and 2, based on the PSEs and SDs of D2, D3 and D4 listed in Table 1. The linear averaging model estimates were calculated by directly averaging the PSEs of D2, D3 and D4 listed in Table 1. (B) Averages of all eleven participants. Both precision weighted model estimates and linear averaging model estimates are broadly consistent with the measured TR results. The errorbars indicate the standard error of the means.

## DISCUSSION

Here we demonstrated that when adapting to thermal referral illusion, the aftereffect depended on the spatial configuration of the fingers in the test phase. When asking about the thermal perception of the whole hand (3-finger) participants’ responses were consistent with having adapted to the illusory rather than physical stimulation. In contrast, when asking about the individual finger perception, participants’ reports were more like having adapted to the physical values. Although the pattern of our results share some phenomenological similarities with contingent aftereffects found in other sensory domains (McCollough, 1965; Walker et al., 1981; Roseboom and Arnold, 2011), the implication that physiological zero would depend on the contingency of finger configuration in recent sensory history is highly speculative as the resetting of the physiological zero occurs at the peripheral level (Fujita et al., 2013). Therefore, our results likely better reflect a process involving only peripheral adaptation. Following peripheral adaptation, such that each finger was adapted to a new physiological zero, presenting the 3-finger configuration in the test phase with a uniformly thermal test display would lead to different apparent temperatures across the 3 fingers. If the sum of these thermal sensations is similar to 35oC, we would find a matched PSE (see Test 2, see Figure 4). The apparent temperatures across 3 fingers under this situation can in fact be estimated based on our data, which showed that the physiological zeros of D2, D3 and D4 after adapting to TR stimulation were [35.3 33.5 35.3] oC, if we assume D4 would behave similarly as D2 (see Test 4 and Test 7 in Figure 4). The shift in apparent temperature as a result of the resetting of physiological zero can be estimated by the difference between the reference temperature, 35oC, and the physiological zero of an adaptation pattern. This follows that the shift in apparent temperature of the three fingers would be [-0.3 1.5 −0.3] oC. Based on this estimation, touching a test display of [35 35 35] oC after TR adaptation would give apparent temperatures of [34.7, 36.5, 34.7] oC across 3 fingers. In light of the results of experiment 2 that the temperature integration across the 3 fingers follows of a simple averaging scheme, the overall apparent temperature would be 35.3 oC, which is indistinguishable from the reference temperature of 35 oC as the precision of temperature estimation is larger than 1oC (see Table 1). Consequently, the whole hand adaptation effect (Test 2, Fig. 4) would be consistent with a peripheral only adaptation. In short, our results support that adaptation takes place only at a peripheral stage where information about temperature inputs are preserved for each finger, and that the TR occurs after this stage.

Our results show that thermal adaptation precedes thermal-tactile integration. Regarding the sensory processing hierarchy for thermal touch, the question remains is in how centrally in this thermo-tactile processing stream does the interaction that produces TR occur? It is known that tactile and thermal sensory systems have physiologically separate ascending sensory pathways (Darian-Smith, 1984) and that their representations in the brain occupy different cortical areas: Discriminative tactile sensations are mediated by the somatosensory cortex, whereas the haptic capacity of thermal sensations is subserved by the dorsal posterior insular cortex (Craig et al., 2000) and/or parietal–opercular (SII) cortex (Mano et al., 2017).

Convergence of thermal and tactile inputs at the subcortical level does exist, but TR is unlikely to occur at the spinal level as all the temperature specific neurons found in the superficial laminae of spinal dorsal horn, where thermoreceptors exclusively terminate, are insensitive to mechanical inputs (Craig and Hunsley, 1991; Dostrovsky and Craig, 1996). Moving further up the sensory stream is the thalamus, a relay center for all incoming sensory inputs. The thalamic relay site for temperature sensation is the posterior part of the thalamic ventromedial nucleus (VMpo), which receives a dense, topographic input from the temperature specific spinothalamic lamina I neurons (Craig et al., 1994; Davis et al., 1999). VMpo is medial and ventroposterior to the thalamic tactile relay nucleus, the ventrocaudal nucleus (VC). There, some neurons have been found to be sensitive to both mechanical and cooling stimuli (Lenz and Dougherty, 1998). However, stimulation at the sites in VC did not elicit cold sensations. Thus, it is not clear whether these multimodal neurons are involved in mediating the sensation of touch or temperature and whether they are related to the thermo-tactile interactions of TR. In brief, these neurophysiological findings suggest that TR, referral of thermal sensations to sites of tactile stimulation, is not merely a hard-wired process that involves subcortical thermoceptive pathway operations. Rather, TR likely results from crossmodal processing at cortical level.

A possible underlying mechanism for TR is that it is driven by high-level crossmodal interaction, and at the same time, subject to constraints posed by low-level organization of the thermoceptive pathway. Thermal sensations are mediated by the small-fiber spinothalamic system, and the neurons on which the spinothalamic fibers terminate have huge receptive fields (Rose and Mountcastle, 1959; Mountcastle, 1961; Han et al., 1998; Craig et al., 2000; Bowsher, 2005). A thermally neutral site might also be activated if its receptive field overlaps with those being physically stimulated. This diffuse nature and the tendency that the thermal sense summates spatially separate inputs (Cain, 1973; Marks and Stevens, 1973) would influence the temperature estimate of each site when multiple sites are in contact. This can be seen in our data that new physiological zeros of the TR adaptation (Test 4 and 7, Figure 4) were not identical to those of the physical temperatures (Test 5 and 8, Figure 4): The PSE of D3 was greater and the PSE of D2 was smaller than those of the corresponding physical temperatures. The constraint of receptive field overlapping explains the effect that TR can be diminished by increasing somatotopic distance, rather than spatial distance, among the stimulated sites (Green, 1978; Ho et al., 2010) and that referral of thermal sensations could also occur without touch (Cataldo et al., 2016). For the crossmodal interactions at the cortical level, in a previous paper (Ho et al., 2011) we proposed that thermal referral is mediated by two separate processes: one determines apparent intensity and the areal extent of the thermal stimulation (how much); the other determines localization of the resulting sensation from the apparent sites of tactile stimulation (where). In the present study, we further demonstrated that the temperature estimation for the 3 fingers under TR (how much) followed an averaging scheme, consistent with Bayesian combination of thermal input from each finger. The integration of the how much (temperature estimation) and where (localization) processes gives a cross-modal estimation that is coherent across thermal and tactile inputs and uniform across all the points in contact.

Our findings support the idea that the combination of different features in thermal-tactile perception follows inference processes for the purpose of coherent object perception. Previous work has shown that inferences regarding visual properties change haptic estimates, such as temperature, weight, surface texture and size (Lederman et al., 1986; Ernst and Banks, 2002; Brayanov and Smith, 2010; Ho et al., 2014). TR similarly appears to be driven by a mechanism in which inferences regarding tactile properties change temperature estimates. That is, the tactile modality signals a homogeneous surface, so the inference for the cause of the spatially incoherent thermal and tactile inputs is in favor of the hand touching a surface of a single, spatially coherent object, rather than a surface with different temperatures at sites in contact. Given that the perceived temperature of objects in contact with the skin is the key to material recognition (e.g. coldness of metal and the warmness of fabrics) (Jones, 2016; Ho, 2018), this kind of crossmodal inference enables us to obtain the information of the material properties without being distracted by small local variations and facilitate object perception in complicated and noisy natural environments (Kersten et al., 2004).

The estimation of tactual temperature is a highly flexible process. When a surface is spatially homogenous, the veridical estimate can be obtained. However, when confronted with a surface with heterogeneous thermal inputs, such as the TR stimulation, the process could involve inference processes related to object perception. Thermal grill illusion (TGI) is another example that demonstrates the flexibility of tactual temperature estimation (Craig and Bushnell, 1994; Green, 2002). In contrast to TR, thermal grill illusion relates more to the estimation of bodily status (Marotta et al., 2015), which presumably involves inferences about the physiological state of the body (Craig, 2002).

In conclusion, we have demonstrated that thermal adaptation occurs prior to thermo-tactile integration in TR. We further showed that the temperature integration under TR results from averaging across fingers, consistent with Bayesian cue combination processes found in other sensory dimensions. These findings support the idea that the interaction of thermal and tactile systems follows common multisensory inference processes likely in service of resolving inconsistent information from thermal and tactile modalities across different points of contact. Such processes are key our ability to manually explore and identify objects based on their shape and material.

